# The Nuclear Pore Complex Facilitates Centriole-Nuclear Attachment in Spermatids

**DOI:** 10.64898/2026.04.28.721503

**Authors:** Danielle B. Buglak, Brian J. Galletta, Nasser M. Rusan

**Affiliations:** Cell and Developmental Biology Center, National Heart Lung and Blood Institute, National Institutes of Health, Bethesda, MD

## Abstract

Proper connection between the sperm head and tail is critical for fertility and is mediated by the head-tail coupling apparatus (HTCA). Recent evidence suggests that the nuclear pore complex (NPC) may be important in male fertility, though a specific role at the HTCA has not been described. To investigate this, we performed a testis-specific RNAi screen targeting nucleoporins of the NPC. We identified Nup133 and Nup107 as regulators of HTCA development. We found that Nup133 and Nup107 were required to form the initial connection between the nucleus and centriole during HTCA establishment. We determined that failure to build the HTCA following Nup133 and Nup107 depletion was due to loss of nuclear envelope dynein/dynactin. Finally, we showed that loss of the NPC cytoplasmic filament component Nup358 results in the most severe centriole detachment phenotype, thus potentially functioning as the dynein anchor. Together, our data indicate that NPCs are critical regulators of early HTCA establishment and are required to recruit dynein to the nuclear envelope to bring the nucleus and centriole together during spermiogenesis.

## INTRODUCTION

Infertility is common, affecting around 15% of couples, with male infertility accounting for approximately half of these cases (Babakhanzadeh et al., 2020; Kumar and Singh, 2015; Pacheco et al., 2023). Male fertility depends on proper development of sperm, which are comprised of a head containing the genetic material, a tail (flagellum) that provides force for swimming, and a sperm neck that connects the head and tail, and contains a centriole or basal body (**Figure 1A**). The proteins and structures within the neck that directly facilitate the connection between the head and tail are known as the head-tail coupling apparatus, or HTCA (**Figure 1A**, Gene-Ontology-Term #0120212). Failures at the HTCA result in a severe type of infertility known as acephalic spermatozoa syndrome, which impacts species across the animal kingdom (Anderson et al., 2009; Baccetti et al., 1989; Blom and Birch-Andersen, 1970; Chemes et al., 1987; Chemes et al., 1999; Deng et al., 2022; Kracklauer et al., 2010; Lai et al., 2016; Li et al., 2004; Nie et al., 2022; Shang et al., 2018; Shang et al., 2017; Sitaram et al., 2012; Texada et al., 2008; Wang et al., 2022; Xia et al., 2024; Zhang et al., 2021a; Zhang et al., 2021b; Zhu et al., 2018; Zhu et al., 2016).

**Figure 1:**
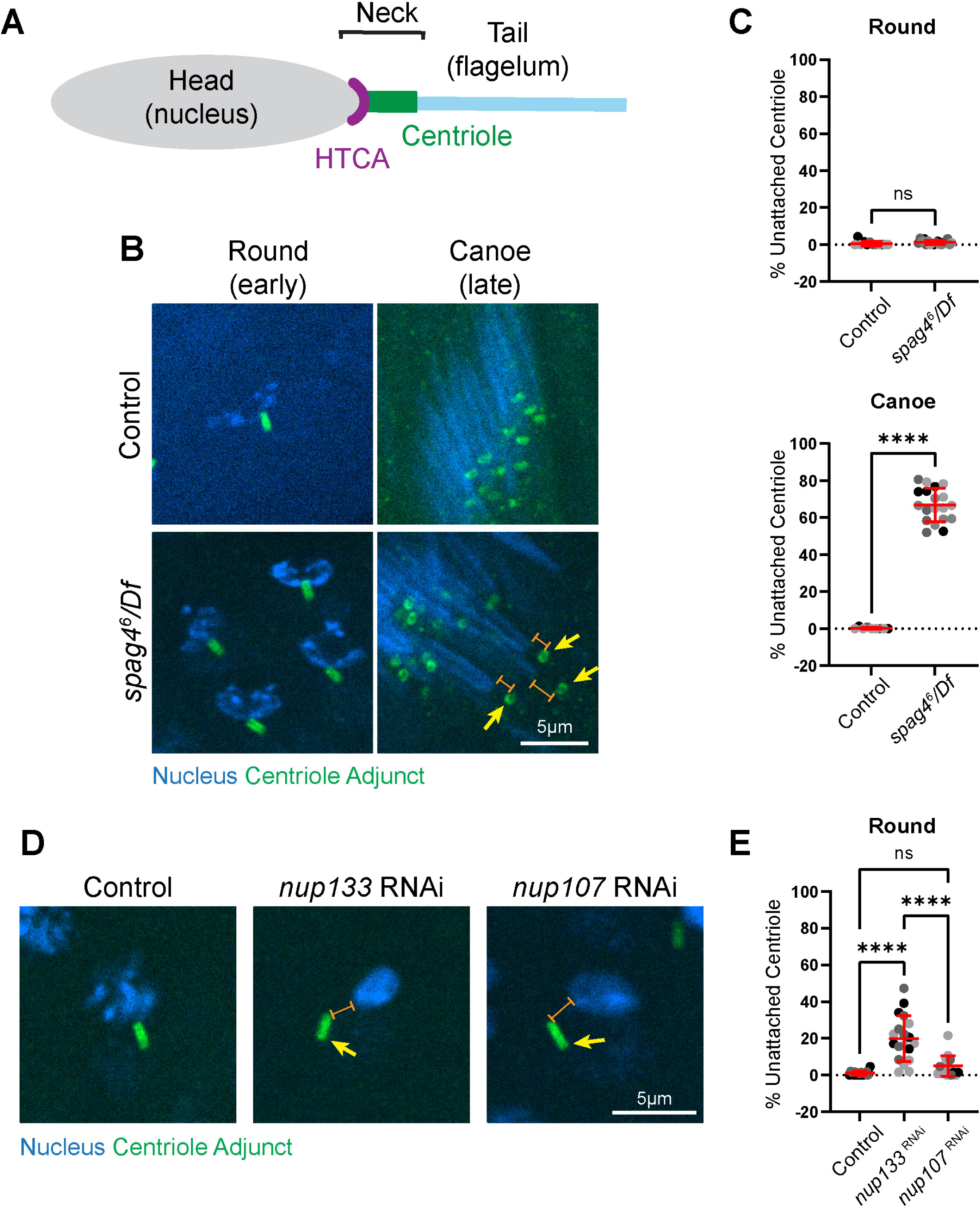
Nup133 and Nup107 function early at the HTCA, while Spag4 functions at later stages. **(A)** Cartoon depicting the sperm HTCA. The head (nucleus, gray) connects to the centriole (green) via the HTCA (purple). The sperm neck encompasses the entire region between the head and tail. **(B)** Representative images of Round and Canoe spermatids of the indicated genotypes. Blue, nucleus (DAPI); green, centriole adjunct (Asl). Yellow arrow indicates detached centriole. Orange brackets show separation between nucleus and centriole. Scale bar 5um. **(C)** Quantification of HTCA attachment in Round and Canoe spermatids for control (*yw*, Round: n=12 ROI, 9 testes; Canoe: n=9 ROI, 7 testes) and *spag4^6^/Df* (*spag4^6^/Df(2L)BSC244,* Round: n=18 ROI, 13 testes; Canoe: n=19 ROI, 13 testes). **(D)** Representative images of Round spermatids of the indicated genotypes. Blue, nucleus (DAPI); green, centriole adjunct (Asl). Yellow arrow indicates detached centriole. Orange bracket shows separation between nucleus and centriole. Scale bar 5um. **(E)** Quantification of HTCA attachment in Round spermatids for control (*Bam-Gal4/+*, n=19 ROI, 16 testes), *nup133* RNAi (*nup133 RNAi/+; Bam-Gal4/+,* n=19 ROI, 17 testes), and *nup107* RNAi (*nup107 RNAi/+; Bam-Gal4/+,* n=15 ROI, 12 testes). ns=not significant; ****=*p*≤0.0001.

The HTCA is fundamentally a linkage between the nucleus and centriole, and is composed of nuclear envelope, cytoplasmic, and centriole components (Buglak et al., 2025). Development of the HTCA begins immediately after meiosis in a process called HTCA Establishment (Buglak et al., 2025; Buglak et al., 2024; Galletta et al., 2020), which occurs in two subphases – Search and Attachment. During the ‘Search’ phase in *Drosophila*, dynein tethered at the nuclear envelope interacts with microtubules nucleated at the proximal end of the centriole to bring the nucleus and centriole together (Anderson et al., 2009; Galletta et al., 2020; Li et al., 2004; Sitaram et al., 2012). During the ‘Attachment’ phase the initial connection at the HTCA occurs. Thus, one key step in forming the HTCA is proper localization of dynein to the nuclear envelope, though the precise mechanism of anchoring dynein is unknown.

Following its establishment, the HTCA is remodeled as part of a process called HTCA Maintenance and Remodeling (Buglak et al., 2025; Buglak et al., 2024). Here too, the precise HTCA molecular linkage is unknown, though some critical components have been discovered, such as the inner-nuclear-membrane protein SUN5 and the cytosolic proteins CENTLEIN and PMFBP1 in mammals (Deng et al., 2022; Lu et al., 2021; Nie et al., 2022; Shang et al., 2018; Shang et al., 2017; Xia et al., 2024; Zhang et al., 2021a; Zhang et al., 2021b; Zhu et al., 2018; Zhu et al., 2016). In *Drosophila*, Spag4, the ortholog of mammalian SUN5, and the cytosolic protein Yuri Gagarin (Yuri) are critical for HTCA function (Buglak et al., 2024; Kracklauer et al., 2010; Moecking et al., 2026; Texada et al., 2008). Undoubtedly, there are critical components of both HTCA establishment and maintenance yet to be identified.

One possible contributor to the HTCA is the nuclear pore complex (NPC), which is a large macromolecular complex in the nuclear envelope important for bidirectional transport of macromolecules. Additionally, the NPC can play transport-independent roles in the nuclear envelope. During mitosis, the NPC is critical for positioning centrosomes at the nuclear envelope during prophase via interactions with microtubule motors (Bolhy et al., 2011; Splinter et al., 2010). Recently, there has been some evidence that the NPC is important in male fertility. A patient with unexplained infertility harbored a variant in Nup210L (Arafah et al., 2021), and mutations in NUP210L and BAF-L, a spermatid-specific paralogue, also cause infertility in mice (Al Dala Ali et al., 2026; Al Dala Ali et al., 2024; Walters et al., 2009). Furthermore, disrupted nucleoporins have been described in patients and mice with infertility (Bragina et al., 2024; Potgieter et al., 2023), and the NPC is dramatically remodeled in mature human sperm (dos Santos et al., 2024), suggesting that regulation of NPCs is important during sperm development.

In *Drosophila*, NPCs have also been linked to sperm development, where Nup358 links the nucleus and a bundle of microtubules known as the manchette (also referred to in flies as the dense complex), which is necessary for nuclear reshaping (Li et al., 2023). A specific role for the NPC at the HTCA has not been described, but a recent study in mice showed that Nup93 interacts with the HTCA protein SUN5 (He et al., 2024), opening the possibility that the NPC could be an important regulator of HTCA establishment or maintenance. Here, we sought to uncover a role for the NPC at the HTCA using a testis-specific RNAi screen of nucleoporins in *Drosophila*. We identified three nucleoporins, Nup133, Nup107, and Nup358 as important regulators of early HTCA establishment that recruit dynein to the nuclear envelope in spermatids.

## RESULTS & DISCUSSION

### NPC proteins are necessary for HTCA establishment and fertility

Our previous work on the inner nuclear membrane SUN-domain protein Spag4 revealed that it undergoes significant remodeling during sperm development (Buglak et al., 2024; Kracklauer et al., 2010). Specifically, Spag4 transitions from encircling the nucleus to a polarized crescent towards the centriole in Round spermatids, finally accumulating and remodeling at the centriole-nucleus contact site (Buglak et al., 2024; Kracklauer et al., 2010). This highly dynamic localization suggested that Spag4 functions at multiple stages of sperm development. We therefore examined the entire process of centriole-nuclear attachment in *spag4* mutants and discovered that *spag4* null spermatids achieved normal centriole-nuclear attachment in early spermatids, but lost attachment in later stages (**Figure 1B,C**). This indicates that Spag4 mainly functions in HTCA maintenance and that a separate, unidentified mechanism must exist for the initial “capture” of the centriole during HTCA establishment as cells exit meiosis.

Given the known role for NPCs in positioning centrosomes in prophase of dividing cells (Bolhy et al., 2011; Splinter et al., 2010), we tested the hypothesis that NPCs play a role in HTCA establishment. We performed a loss-of-function screen of 27 nucleoporin proteins and assessed the ability of centrioles to dock to the nuclear envelope (**Table S1**). We used a germline specific Gal4-driver to drive UAS-RNAi against each of the 27 proteins to identify cell autonomous effects within the germline and avoid pleiotropic effects from more global knockdowns and mutations. We observed several phenotypes, including abnormalities in nuclear shaping and defects that arise from meiotic errors such as atypical centriole numbers and oversized mitochondrial derivatives with multiple nuclei and centrioles in spermatids (**Table S1**, **Figure S1**). Several of the tested nucleoporin knockdowns exhibited HTCA phenotypes, and interestingly, the developmental timing of these defects differed. We classified these HTCA phenotypes as “early” if they appeared in Round spermatids or “late” if they appeared in Leaf or Canoe spermatids (**Table S1**, **Figure S1**). We found that using two independent RNAi lines for each of Nup133 and Nup107 resulted in reduced centriole-nuclear attachment in Round spermatids as well as nuclear shaping defects (**Table S1, Figure S1**), suggesting they could play a role upstream of Spag4 in early HTCA establishment. Nup133 and Nup107 are both part of the NPC Y-complex, a group of nucleoporins that form the outer structural rings of the NPC (Harel et al., 2003; Kelley et al., 2015; Vollmer et al., 2015; Walther et al., 2003). Our follow-up quantitative analysis in spermatids stained for the centriole adjunct (as a proxy for the centriole) revealed that 20% of centrioles in *nup133* knockdown cells and 5% in *nup107* knockdown cells are unattached to the nucleus in Round spermatids (**Figure 1D,E**).

A lack of nucleus-centriole attachment in Round spermatids is typically the result of errors with HTCA establishment and frequently an inability of the nucleus and centriole to come together during the nuclear search phase, as is the case with dynein mutants (Anderson et al., 2009; Li et al., 2004; Sitaram et al., 2012). To investigate the dynamics of nuclear search following Nup depletion, we performed live cell imaging of whole mount testes expressing the nuclear marker H2A::RFP and centriole marker PACT::GFP in both control and RNAi conditions. We found that in control cells exiting meiosis, the nucleus and centriole moved closer together before forming an initial attachment (**Figure 2A,B**). However, in *nup133* and *nup107* RNAi spermatids, the nucleus and centriole did not efficiently form attachments. Both the *nup133* and *nup107* RNAi spermatids resulted in fewer than 50% of centrioles positioned within 0.5um from the nucleus by 45 min post-meiosis (compared to 75% in controls, **Figure 2B**) and fewer centrioles in the knockdown conditions reaching the nuclear surface by 70 minutes post-meiosis (**Figure 2B**, y=zero) in agreement with our fixed analysis (**Figure 1E**). Collectively, these data suggest that Nup133 and Nup107 are required to properly position the centriole during nuclear search and attachment, providing further evidence that the NPC is important in early HTCA establishment.

**Figure 2:**
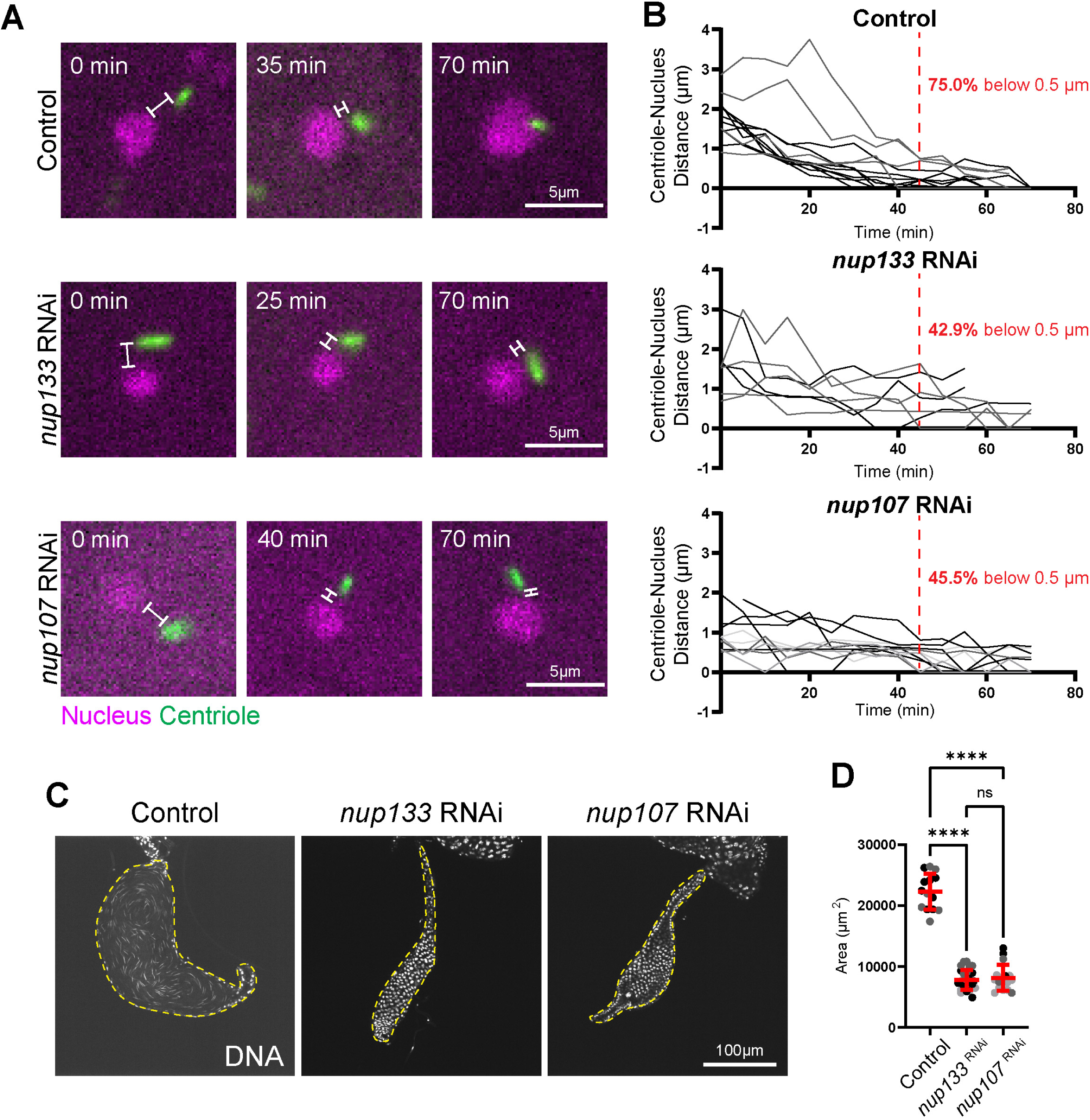
Nup133 and Nup107 are necessary for HTCA establishment and fertility. **(A)** Stills from live imaging of HTCA attachment for indicated genotypes. Magenta, nucleus (H2A::RFP); green, centriole (PACT::GFP). White brackets show separation between nucleus and centriole. Scale bar 5um. **(B)** Quantification of distance between the nucleus and centriole over time in controls (*H2A::RFP; PACT::GFP,* n=12 spermatids, 2 testes), *nup133* RNAi (*H2A::RFP/nup133 RNAi; PACT::GFP/Bam-Gal4,* n=7 spermatids, 2 testes), and *nup107* RNAi (*H2A::RFP/nup107 RNAi; PACT::GFP/Bam-Gal4,* n=11 spermatids, 4 testes). Red dotted line represents the 45 min mark. Number of spermatids with a nucleus-centriole distance less than 0.5um at this time point are shown in red for each genotype. **(C)** Representative images of seminal vesicles of the indicated RNAi knockdowns stained with DAPI (DNA). Yellow dashed outline shows seminal vesicles. **(D)** Quantification of seminal vesicle area for controls (*Bam-Gal4/+,* n=14 seminal vesicles), *nup133* RNAi (*nup133 RNAi/+; Bam-Gal4/+,* n=28 seminal vesicles), and *nup107* RNAi (*nup107 RNAi/+; Bam-Gal4/+,* n=17 seminal vesicles). ns=not significant; ****=*p*≤0.0001.

Finally, as an endpoint assay, we investigated if Nup133 and Nup107 were required for male fertility. We examined the seminal vesicles following *nup133* and *nup107* RNAi knockdown and found they were significantly smaller and devoid of sperm (**Figure 2C,D**), indicating that Nup133 and Nup107 are critical for fully functional sperm development. This complete sterility is likely a synergistic result of early HTCA failure combined with the later nuclear shaping/manchette defects.

Collectively, our data suggest a two-step model for HTCA formation, where Nups and Spag4 perform a sequential molecular handoff over the course of spermiogenesis. We propose that the NPC functions as the initial “capture” machinery during the dynamic search phase, potentially repurposing the mitotic prophase tethering mechanism (Bolhy et al., 2011), while Spag4 functions downstream to permanently “lock” the centriole in place.

### Nup133 and Nup107 localize and remodel in conjunction with the HTCA

A lack of centriole-nuclear connection following Nup133 or Nup107 depletion suggested that these proteins likely localize to the HTCA. To investigate their subcellular localization we used UAS-GFP::Nup133 (driven by *tubulin-Gal4*) or GFP::Nup107 (driven by its own promoter) and found that they both localize to the nuclear envelope in spermatocytes then rapidly form a polarized crescent towards the HTCA in Round spermatids (**Figure 3A,B**), consistent with other studies examining the localization of Nup358, Nup50, and NDC1 at the HTCA in rodents and flies (Fan et al., 1997; Lai et al., 2016; Li et al., 2023). To further confirm localization of the NPC in Round spermatids, we used transmission electron microscopy and found that NPCs were polarized towards the HTCA (**Figure 3C, yellow arrows**), consistent with other EM studies (Ho, 2010).

**Figure 3:**
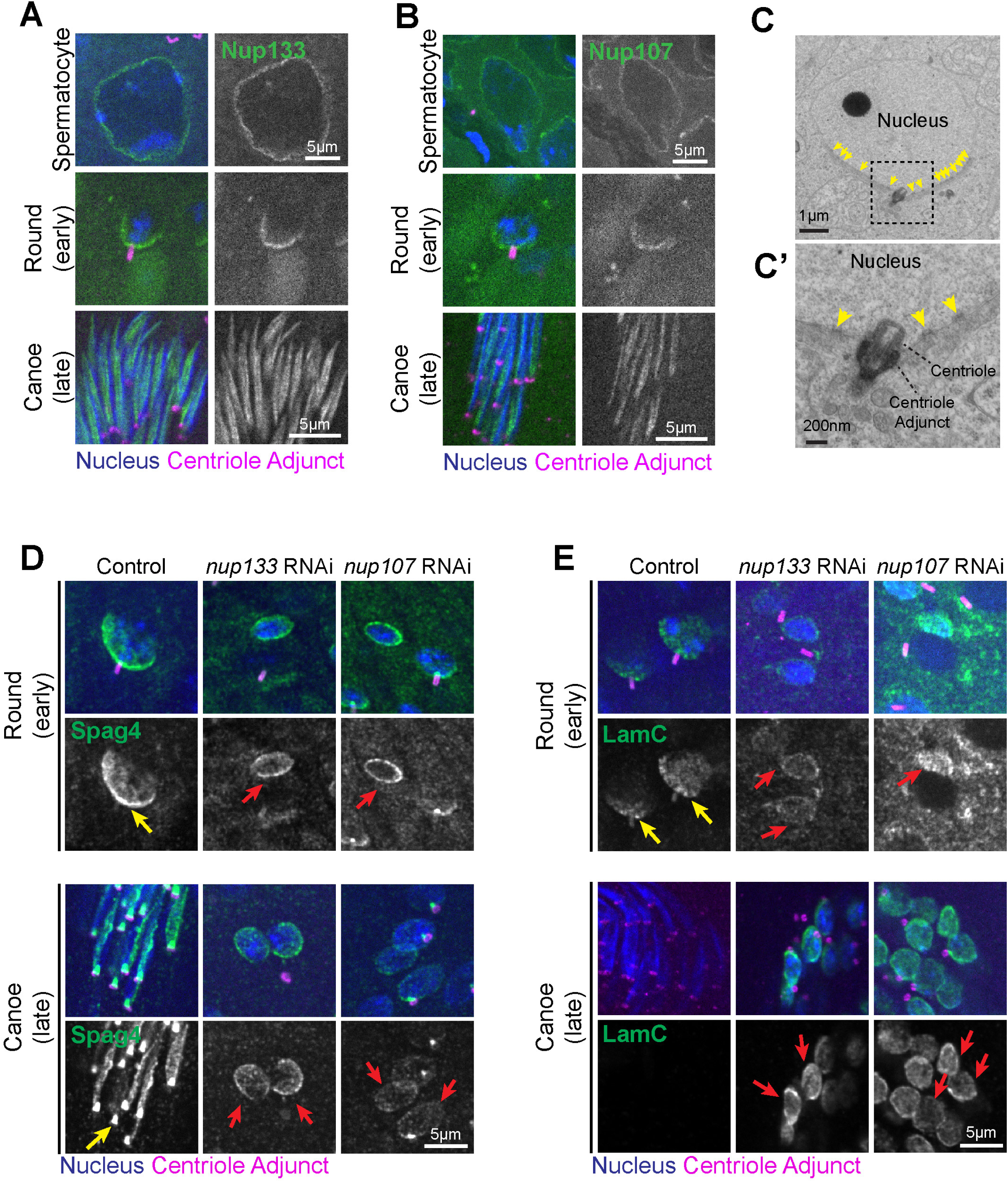
Nup133 and Nup107 are needed for proper localization of HTCA and nuclear envelope proteins. **(A)** Representative images of Nup133 (UAS-GFP::Nup133, green) localization in wild-type Spermatocytes, Round spermatids, and Canoe spermatids. Blue, nucleus (DAPI); magenta, centriole adjunct (Asl). **(B)** Representative images of Nup107 (GFP::Nup107, green) localization in wild-type Spermatocytes, Round spermatids, and Canoe spermatids. Blue, nucleus (DAPI); magenta, centriole adjunct (Asl). Scale bar 5um. **(C-C’)** Electron micrograph of a wild-type spermatid showing nuclear pores (yellow arrows) clustered towards the HTCA. Scale bar 1um **(C),** 200nm **(C’)**. **(D-E)** Representative images of Round and Canoe spermatids for the indicated genotypes. Blue, nucleus (DAPI); magenta, centriole adjunct (Asl); green, Spag4 (Spag4::6myc) **(D)** or LamC **(E)**. Yellow arrows indicate normal polarized Spag4 or LamC localization. Red arrows indicate abnormal Spag4 or LamC surrounding the nucleus. Scale bars 5um.

Interestingly, Nup133 and Nup107 localization dramatically changed in later stage spermatids (Canoe), localizing along one side of the nucleus towards the manchette (**Figure 3A,B**). This is consistent with the NPC’s role in linking the manchette to the nucleus (Li et al., 2023), and likely explains the nuclear reshaping defect that was also observed following *nup133* or *nup107* RNAi (**Figure S1; Table S1**). The migration of Nups away from the HTCA and towards the manchette provides strong support for our “hand-off” model to repurpose the Nups from capturing the centriole to sperm head reshaping, which further supports the model that Nup133 and Nup107 are not required for later HTCA maintenance.

### Nup133 and Nup107 are required for localization of HTCA components

Because we observed failed HTCA establishment in spermatids following *nup133* or *nup107* RNAi, we next tested whether loss of NPC proteins impacts the localization of other key HTCA and nuclear envelope proteins. An important step in HTCA development is the transition of nuclear envelope proteins, such as Spag4 and the nuclear lamina, from completely encircling the nucleus to a polarized crescent facing the centriole (Buglak et al., 2024; Fabbretti et al., 2016; Kracklauer et al., 2010). We found that Spag4 was unable to properly polarize in *nup133* or *nup107* RNAi flies, and instead encircled the nucleus (**Figure 3D, red arrows**). This uniform distribution of Spag4 persisted into Canoe spermatids, which suggests the Nup-dependent interaction of the nucleus with centriolar microtubules acts as the symmetry-breaking event that cues the subsequent polarization of Spag4. Without the Nup anchor, the nuclear envelope remains radially symmetric, unable to define the distinct HTCA face.

Additionally, LamC was not only unable to polarize, but was also retained at the nuclear envelope in Canoe *nup133* and *nup107* RNAi spermatids (**Figure 3E**). We suspect that the retention of LamC contributes to the impaired nuclear reshaping in Nup-depleted spermatids. How depletion of NPC components leads to lamin retention is not clear and will be important for further study.

### Protein transport across the nuclear envelope is largely preserved upon Nup133 and Nup107 depletion

Because the NPC is critical for nuclear transport, we considered the possibility that loss of Nup133 or Nup107 prevented normal nuclear transport, thus indirectly impacting localization of HTCA proteins and resulting in the observed defects in HTCA establishment. For example, interference with Nup107 or Nup133 function in other systems results in mRNA export defects, but does not always affect protein import (Boehmer et al., 2003; Doye et al., 1994; Vasu et al., 2001; Walther et al., 2003). We found that a transgenic NLS-tagged GFP construct was able to enter the nucleus in Nup-depleted spermatids, though accumulation in the nucleus after meiosis took longer than controls (**Figure S2A**). Furthermore, Protamine B, whose mRNA is translationally repressed in the cytoplasm prior to its transport into the nucleus of spermatids (Jayaramaiah Raja and Renkawitz-Pohl, 2005), accumulated in the nuclei of late canoe spermatids similar to controls (**Figure S2B**).

Finally, we analyzed Asunder, a critical upstream regulator of dynein necessary for proper HTCA development and perinuclear dynein localization (Anderson et al., 2009; Sitaram et al., 2012). In both *Drosophila* and mammals (via its highly conserved ortholog ASUN/INTS13), Asunder promotes dynein enrichment at the nuclear envelope (Jodoin et al., 2012; Jodoin et al., 2013); however, it is initially localized to the nucleoplasm before being exported to the cytoplasm to properly function (Anderson et al., 2009; Jodoin et al., 2013; Sitaram et al., 2012). We considered the possibility that Nup depletion might trap Asunder in the nucleus, thereby indirectly causing the observed failure in dynein recruitment. However, we found that the export of Asunder into the cytoplasm and its subsequent exclusion from the nucleus are unaffected in spermatids following *nup133* or *nup107* RNAi (**Figure S2C**). This demonstrates that our observed loss of dynein at the nuclear envelope following NPC disruption is not merely a downstream consequence of sequestered Asunder, but rather operates via a distinct, Asunder-independent mechanism.

While we cannot entirely rule out a nuclear transport defect, collectively these findings indicate that transit across the nuclear envelope remains intact, and that the centriole attachment defects in *nup133* or *nup107* RNAi are likely a result of losing a transport-independent physical interaction of HTCA machinery with the NPC.

### NPCs are required to recruit dynein-associated machinery

During early HTCA establishment, dynein at the nuclear membrane interacts with microtubules emanating from the centriole to reel the two organelles together to establish the HTCA (Anderson et al., 2009; Galletta et al., 2020; Li et al., 2004; Sitaram et al., 2012). While previous work suggests that dynein organization in early spermatids somehow depends on Spag4 (Kracklauer et al., 2010), it is not known what anchors dynein to the nuclear envelope at this stage. We hypothesized that NPCs are the dynein anchor based on the negative effects of *nup133* or *nup107* RNAi on centriole attachment, in addition to the failed polarization of Spag4 and LamC. Indeed, we found that both the dynactin subunit 3 (DCTN3, p24) and the dynein adaptor Lis-1 had significantly reduced localization to the nuclear envelope in *nup133* or *nup107* RNAi Round spermatids (**Figure 4A-D**).

**Figure 4:**
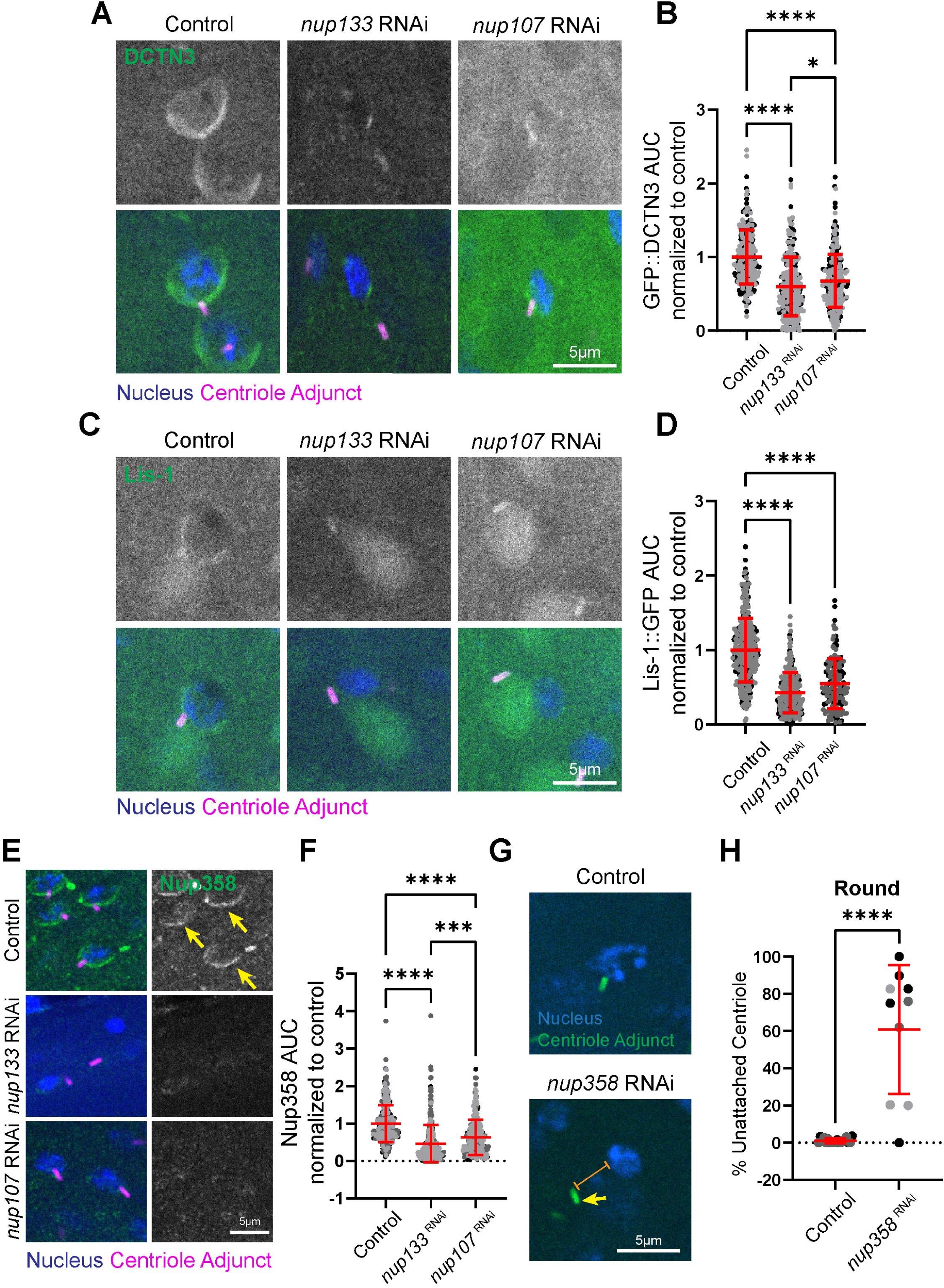
Loss of Nup133 and Nup107 are required for dynactin, Lis-1, and Nup358 recruitment to the nuclear envelope. **(A)** Representative images of Round spermatids of the indicated genotypes. Blue, nucleus (DAPI); magenta, centriole adjunct (Asl); green, DCTN3 (GFP::DCTN3). Scale bar 5um. **(B)** Quantification of DCTN3 at the nuclear envelope in Round spermatids for controls (*spag4::6myc/IF; ubi-GFP::DCTN3/Bam-Gal4,* n=291 spermatids, 17 testes), *nup133* RNAi (*spag4::6myc/nup133 RNAi; ubi-GFP::DCTN3/Bam-Gal4,* n=324 spermatids, 15 testes), and *nup107* RNAi (*spag4::6myc/nup107 RNAi; ubi-GFP::DCTN3/Bam-Gal4,* n=311 spermatids, 12 testes). **(C)** Representative images of Round spermatids of the indicated genotypes. Blue, nucleus (DAPI); green, centriole adjunct (Asl); magenta, Lis-1 (Lis-1::GFP). **(D)** Quantification of Lis-1 at the nuclear envelope in Round spermatids for controls (*spag4::6myc/CyO* or *IF; Lis-1::GFP/TM3* or *TM6B,* n=404 spermatids, 17 testes), *nup133* RNAi (*spag4::6myc/nup133 RNAi; Lis-1::GFP/Bam-Gal4,* n=260 spermatids, 13 testes), and *nup107* RNAi (*spag4::6myc/nup107 RNAi; Lis-1::GFP/Bam-Gal4,* n=156 spermatids, 7 testes). *\**=*p*≤0.05; ****=*p*≤0.0001. **(E)** Representative images of Round and Canoe spermatids for the indicated genotypes. Blue, nucleus (DAPI); magenta, centriole adjunct (Asl); green, Nup358. Yellow arrows indicate normal polarized Nup358 localization. Scale bars 5um. **(F)** Quantification of Nup358 at the nuclear envelope in Round spermatids for controls (*Bam-Gal4/+*, n=360 spermatids, 19 testes), *nup133* RNAi (*nup133 RNAi/+; Bam-Gal4/+*, n=295 spermatids, 14 testes), and *nup107* RNAi (*nup107 RNAi/+; Bam-Gal4/+*, n=233 spermatids, 11 testes). **(G)** Representative images of Round spermatids of the indicated genotypes. Blue, nucleus (DAPI); green, centriole adjunct (Asl). Scale bar 5um. **(H)** Quantification of HTCA attachment in Round spermatids for control (*Bam-Gal4/+,* n=19 ROI, 16 testes) and *nup358* RNAi (*nup358 RNAi/+; Bam-Gal4/+,* n=10 ROI, 9 testes). *=*p*≤0.05; ***=*p*≤0.001; ****=*p*≤0.0001.

While we provide the first evidence that the NPC are required to recruit dynein to the nuclear envelope in spermiogenesis, NPC-dynein link is not without precedent. For example, the NPC serves as a dynein tether during mitosis via two pathways: 1) a Nup133-CENP-F-NudE/EL pathway and 2) a Nup358-BICD2 pathway (Bolhy et al., 2011; Hu et al., 2013; Splinter et al., 2010). Dynein light chain (DYNLT1) was also shown to interact directly with nucleoporins by yeast 2 hybrid (Sarma and Yaseen, 2013), and most recently, Nups were found to position centromeres in *Drosophila* neuroblasts (Taylor and Cabernard, 2025), likely via a dynein/microtubule-based mechanism. Therefore, it is possible that the spermatid NPC repurposes one of these pathways to tether dynein, or that it uses a unique pathway to link the NPC to the dynein/dynactin complex and its regulators.

### The cytoplasmic filament component Nup358 is required for HTCA establishment

While it is possible that Nup133 directly recruit dynein to the spermatid nuclear envelope using the Nup133-CENP-F-NudE/EL pathway as was shown in prophase cells (Bolhy et al., 2011), it is also possible that the link to dynein is indirect. Given that Nup133 is a member of the Y-complex that provides the structural scaffold for cytoplasmic filament Nups, such as Nup358 (Zhu et al., 2022), we hypothesized that the direct link to dynein could be through the Nup358 pathway (Splinter et al., 2010). To test this, we used *nup133* or *nup107* RNAi and assayed for the localization of Nup358. In contrast to control Round spermatids with clear polarized Nup358 nuclear crescents, loss of *nup133* or *nup107* significantly reduced Nup358 levels such that a crescent was barely detected (**Figure 4E,F**). This suggests that NPC cytoplasmic filaments require Nup133 and Nup107. This result led us to revisit our initial screen, where we had classified Nup358 as having severe meiotic defects (Table S1). A more careful analysis of the *nup358 RNAi* cysts revealed that a low number of spermatids were able to exit meiosis, but displayed significant nucleus-centriole attachment defects (**Figure 4G,H**).

This HTCA establishment defect was more severe than both *nup133* and *nup107* RNAi (**Figure 1E**), suggesting that Nup358 could play a more direct role in dynein recruitment. Also, similar to loss of Nup133 and Nup107, we observed smaller seminal vesicles devoid of sperm following *nup358* RNAi, indicating a critical role in fertility (**Figure S3A,B**).

### Model of NPC-dependent HTCA establishment

Overall, our data supports a model where NPCs function to recruit dynein in a critical window of time during HTCA establishment (**Figure 5A**). During the search phase, the NPC recruits dynein/dynactin and the adaptor Lis-1 to the nuclear envelope where they interact with microtubules to reel the centriole to the nucleus (Anderson et al., 2009; Galletta et al., 2020; Li et al., 2004; Sitaram et al., 2012). NPCs and other nuclear membrane proteins like Spag4 and the nuclear lamina polarize towards the HTCA as dynein brings the nucleus and centriole together (**Figure 5A**; search and attachment). Given that *spag4* mutants did not show a loss of centriole-nucleus attachment during establishment, Spag4 likely does not play a major role in this early process, but appears to depend on the NPC for its polarization towards the HTCA (**Figure 5A,B**).

**Figure 5:**
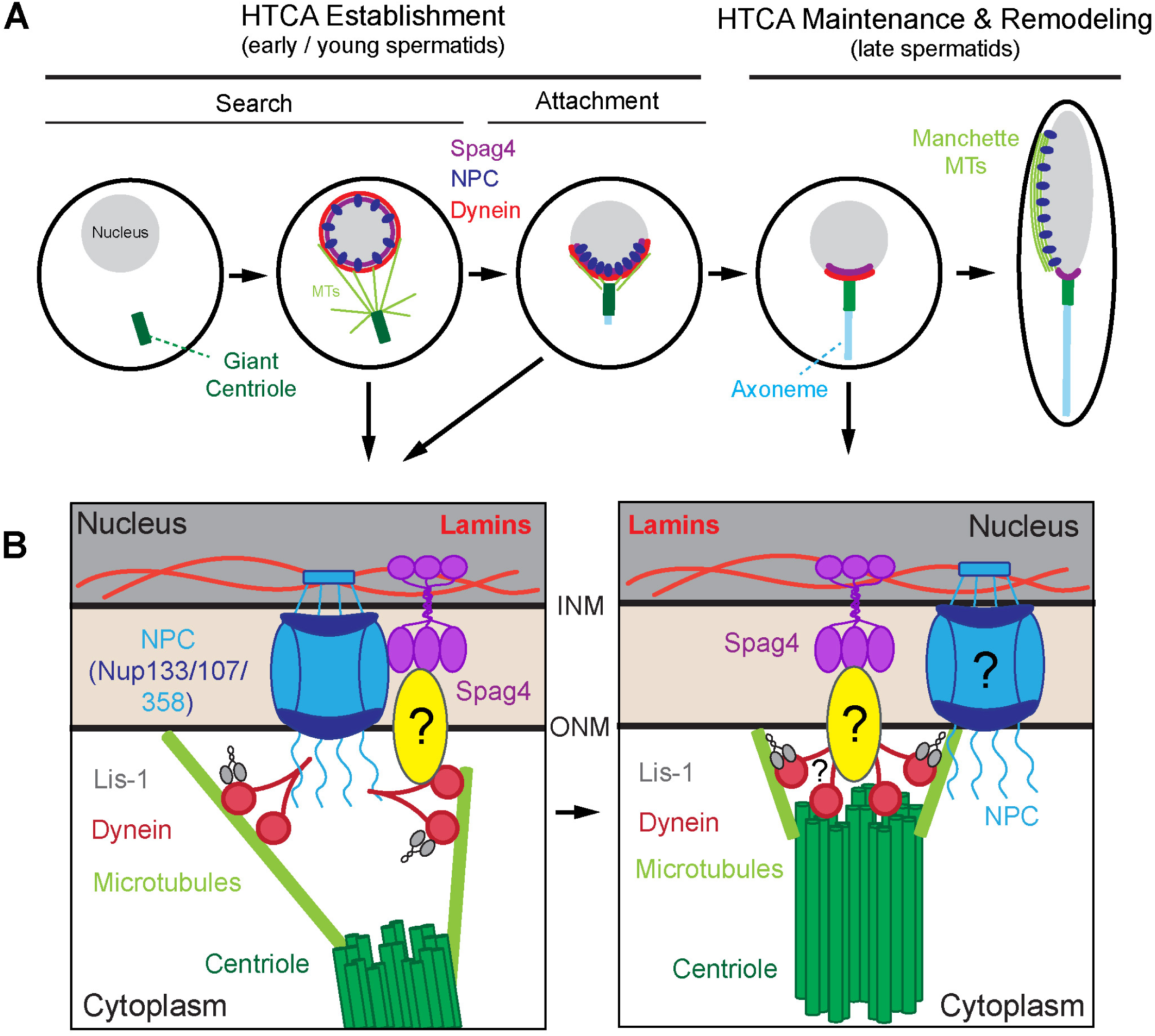
Hand-off model for centriole attachment from NPCs to Spag4. **(A)** HTCA establishment occurs in two phases: Search and Attachment. During search, the nucleus (gray) and centriole (green) come together via dynein (red) which is recruited to the nuclear envelope by the NPC (blue). As the attachment forms, dynein, NPCs, and Spag4 (purple) are polarized towards the HTCA. During HTCA maintenance and remodeling, the NPCs reorganize along one side of the nucleus where they serve to anchor the manchette microtubules (light green) to the nucleus. **(B)** During establishment (left panel), the NPC (via Nup358, Nup133, and Nup107) recruit dynein and the adaptor Lis-1 to the nuclear envelope. These motors then interact with microtubules nucleated from the proximal end of the centriole to bring the nucleus and centriole together. During HTCA maintenance and remodeling (right panel), Spag4 and unknown proteins lock the nucleus and centriole in place. The NPC, dynein, and Lis-1 may play additional roles in the later stages.

During the later stages of HTCA maintenance and remodeling, NPCs relocalize along one side of the nucleus (**Figure 5A**), where they function to link the nucleus and manchette (Li et al., 2023). At this stage, Spag4 takes over as the major contributor to nucleus-centriole attachment, though the specific molecular linkage remains unclear (**Figure 5B**) (Buglak et al., 2025). However, several Nups in our RNAi screen, such as Nup93-2 and Nup43, exhibited late HTCA defects, suggesting that the NPC may indeed play roles during HTCA maintenance and remodeling, possibly functioning alongside Spag4 (**Figure 5B**). Collectively, our work contributes to the growing body of evidence that NPCs are important in male fertility (Al Dala Ali et al., 2024; Arafah et al., 2021; Bragina et al., 2024; dos Santos et al., 2024; He et al., 2024; Li et al., 2023; Potgieter et al., 2023; Walters et al., 2009). While some studies have shown that NPCs are disrupted in infertile individuals (Bragina et al., 2024; Potgieter et al., 2023), the precise contributions of the NPC to sperm development have remained elusive. We demonstrate for the first time that the NPC recruits dynein to the nuclear envelope in spermatids, facilitating HTCA establishment and highlighting the NPC as an important target for studies in infertility.

## Acknowledgements

We thank Samantha L. Smith and the NHLBI Electron Microscopy Core for help with electron microscopy of testes, and Carey J. Fagerstrom for help with cloning. We also thank Rusan lab members as well as Mary Dasso and Alexei Arnaoutov for helpful discussion and reagents. This work was supported by the Division of Intramural Research at the National Heart, Lung, and Blood Institute (ZIAHL006126 to NMR).

## Disclaimer

The contributions of the NIH author(s) are considered Works of the United States Government. The findings and conclusions presented in this paper are those of the author(s) and do not necessarily reflect the views of the NIH or the U.S. Department of Health and Human Services.

## Data availability statement

All relevant data can be found within the article and its supplementary information. The DOI for all raw data is 10.25444/nhlbi.31059577

## MATERIALS AND METHODS

### Flies

*D. melanogaster* were maintained and crosses were performed at 25°C. We used the following strains: *yw, spag4::6myc* (Bloomington Drosophila Stock Center, Bloomington, IN, USA, #29981, #29980), *Bam-Gal4* (Chen and McKearin, 2003), *tubulin-Gal4* (Lee and Luo, 1999), *nup133* RNAi (Bloomington Drosophila Stock Center, Bloomington, IN, USA #58290; Vienna Drosophila Resource Center, Vienna, AUT, #110194), *nup107* RNAi (Vienna Drosophila Resource Center, #22407 and #110759), GFP::Nup107 (Bloomington Drosophila Stock Center, #35514), *H2A::RFP* (Bloomington Drosophila Stock Center, # 23651), *ubi-PACT::GFP* (Galletta et al., 2020; Martinez-Campos et al., 2004), *Lis-1::GFP* (Vienna Drosophila Resource Center, #v318159), *protB::GFP* (Bloomington Drosophila Stock Center, #58406), *spag4^6^* (Bloomington Drosophila Stock Center, #29978), *Df(2L)BSC244/CyO* (Bloomington Drosophila Stock Center, #9718). Fly RNAi lines used for the RNAi screen can be found in **Table S1**. For most experiments, *nup133* RNAi line #58290 and *nup107* RNAi line #110759 were used.

Nup133 cDNA LP09464 from the Gold Collection (DGRC Stock 1668352 ; https://dgrc.bio.indiana.edu//stock/1668352) was used as template for PCR and cloned into pENTR-D/Topo vector. Primers used were: Nup133 F CACCATGGAGAGAAATCTGCAGAAGCAATTATACGGC, Nup133 R GAGTTCCATGTCGTCTGGCTGTTTGAACATGTC. Destination reaction using the Gateway cloning system (ThermoFisher Scientific, Waltham, MA) was performed to move the cDNA segment into the destination vector pPGW (DGRC Stock 1077 ; https://dgrc.bio.indiana.edu//stock/1077 ; RRID:DGRC_1077). DCTN3 cDNA was synthesized as a double stranded linear DNA (Twist Biosciences, South San Francisco, CA) and cloned into pENTR-D-Topo vector. Destination reaction using the Gateway cloning system (ThermoFisher Scientific, Waltham, MA) was performed to move the cDNA segment into the destination vector pUGW (DGRC Stock 1283; https://dgrc.bio.indiana.edu//stock/1283 ; RRID:DGRC_1283). All transgenic flies were generated using standard P-element transformation (Bestgene, Inc.).

### Testis fixation and immunofluorescence

Testes were dissected from 1-6 day old males in Schneider’s medium with antibiotic-antimycotic unless otherwise indicated. Testes were fixed in 4% paraformaldehyde (PFA) for 20 min at room temperature, washed in PBS with 0.3% Triton X-100 (PBST), and blocked in 5% normal goat serum (NGS) in PBST for at least 2h at room temperature. Primary antibody was added overnight in blocking solution at 4°C then washed 3 times for 10 min in PBST. The following primary antibodies were used: mouse anti-9E-10 c-myc (Developmental Studies Hybridoma Bank, Iowa City, IA, USA; Cat# 9E 10-c; 1:1,000), mouse anti-9E-10 c-myc (Thermo Fisher Scientific, Waltham, MA, USA; Cat# MA1-980; 1:3,000), guinea pig anti-Asl (Klebba et al., 2013) (1:10,000), mouse anti-ATP5A (Abcam, Waltham, MA, USA; Cat# Ab14748; 1:500), mouse anti-LamC (Developmental Studies Hybridoma Bank; Cat# LC28.26; 1:100), chicken anti-Nup358 (1:300) (gift from Mary Dasso lab, NICHD). Secondary antibody was added in blocking solution for at least 3h at room temperature. The following secondary antibodies were used: AlexaFluor488 conjugated (Thermo Fisher Scientific; 1:1,000), AlexaFluor568 conjugated (Thermo Fisher Scientific; 1:1,000), AlexaFluor647 (Thermo Fisher Scientific; 1:1,000). DAPI (Thermo Fisher Scientific; Cat# D1306; 1:100) was added alongside secondary antibodies. Testes were then washed 3 times for 10 min in PBST then mounted onto glass slides with AquaPolymount (Polysciences, Inc., Warrington, PA, USA) and a No 1.5 coverslip for confocal imaging.

### Confocal imaging

Confocal images were acquired on a Nikon Eclipse Ti2 (Nikon Instruments, Melville, NY, USA) with a Yokogawa CSU-W1 spinning disk confocal head (Yokogawa Life Science, Sugar Land, TX, USA) equipped with a Prime BSI cMos camera (Teledyne Photometrics, Tucson, AZ, USA) or a Nikon Eclipse Ti-E (Nikon Instruments) with a Yokogawa CSU-X1 spinning disk confocal head (Yokogawa Life Science) equipped with an ORCA-Flash 4.0 CMOS camera (Hamamatsu Photonics) or Kinetix22 sCMOS camera (Teledyne Photometrics). 405, 488, 561, and 641 nm laser lines were used. Spermatids were imaged using a 100xTIRF/1.49 NA oil immersion objective, seminal vesicles were imaged using a 40x/1.30 NA oil immersion objective, and expansion gels were imaged using a 100x/1.35 NA silicone immersion objective. All images were acquired using Nikon Elements software (Nikon Instruments) and analyzed and processed in FIJI (ImageJ, National Institutes of Health, Bethesda, MD, USA).

### Spermatid staging

Round spermatids were staged based on the shape of the nucleus and mitochondria (marked by ATP5A). Spermatids with round mitochondria approximately the same size as the nucleus were classified as “Round.” Leaf and Canoe spermatids were staged as previously described (Buglak et al., 2024). Due to severe nuclear shaping defects in some RNAi experiments, the shape of the centriole adjunct (Asl) was used to assist with staging (Buglak et al., 2024).

### Nucleoporin RNAi screen

RNAi lines for known and predicted *Drosophila* nucleoporins were obtained from the Bloomington Drosophila Stock Center and Vienna Drosophila Resource Center (**Table S1**). Multiple RNAi lines were used where possible. Males carrying the RNAi lines were crossed with virgin females carrying *Bam-gal4* to drive expression of the RNAi in the testes of male progeny. At least 3 testes were imaged for each RNAi line.

Testes were assessed for HTCA defects at each stage of spermiogenesis (Round, Leaf, Canoe, and Needle) by observing the position of the centriole adjunct (marked by Asl) and the nucleus. Separation between the centriole adjunct and nucleus was considered an HTCA defect. HTCA defects appearing in Round spermatids were classified as “early HTCA defects,” while defects that did not appear until Leaf spermatids or later were classified as “late HTCA defects.”

### Centriole attachment quantification

For measurements of Round stages, spermatids with round nuclei and elongated Asl were used. For measurements of centriole attachment in Canoe stages for *spag4^6^* mutants, spermatids were used if the width of the nucleus was <2um. Centrioles touching the nucleus in any capacity were designated as “attached,” while centrioles that did not overlap with the nuclear signal were designated as “unattached.” In some cases, nuclear membrane markers such as LamC, Nup358, and RanGAP were also used to facilitate assessment of attachment. This was tracked using the Cell Counter plugin in FIJI. The percentage of unattached spermatids was then calculated for each image. Due to the challenges of analyzing centriole-nucleus contact in Z, this likely represents an underestimate of the detachment phenotype. Images were only considered if at least two-thirds of a cyst (43 spermatids) could be accurately assessed. For *nup107* and *nup358* RNAi which had meiotic defects, HTCA defects were only assessed in spermatids without obvious meiotic defects (abnormal mitochondria:centriole:nucleus ratio), which may lead to an underestimate of the HTCA phenotype.

### Live imaging

For live imaging, pupal testes were dissected in Schneider’s medium with antibiotic-antimycotic. Testes were mounted on a Lumox dish in dissection medium using Halocarbon oil under a No 1.5 coverslip. Images were acquired on a Nikon Eclipse Ti2 (Nikon Instruments) with a Yokogawa CSU-W1 spinning disk confocal head (Yokogawa Life Science) equipped with a Prime BSI cMos camera (Teledyne Photometrics) using the 488 and 561 nm laser lines and a 40x/1.30 NA oil immersion objective. Images were acquired at room temperature every 5 min for at least 70 min using Nikon Elements software (Nikon Instruments). Movies were processed and analyzed in FIJI (ImageJ). For quantification, spermatids that exited meiosis where the nucleus and centriole had not yet come together were followed for 70 min post-meiosis. A line was drawn between the center of the proximal end of the centriole closest to the nucleus and the edge of the nucleus to determine the distance between the two organelles at each time point. The % of spermatids within 0.5um of the nucleus at 45 min was then determined. For spermatids where a measurement could not be taken at the 45 min frame, the average between the measurements immediately before and after 45 min was used.

### Seminal vesicle area

Seminal vesicles were dissected from virgin males aged for 2 days in the absence of females at room temperature. Seminal vesicles were fixed for 20 min in 4% PFA at room temperature then incubated for at least 3h in DAPI (1:100) in PBST with 5% NGS. The middle Z-slice of each seminal vesicle was used for area calculation. If the entire seminal vesicle could not be visualized in the middle Z-slice, the closest Z-slice to the middle where the entire tissue was visible was used. An ROI was drawn along the outside of the seminal vesicle in FIJI using the DAPI signal as an indicator. The area of this ROI was calculated in FIJI and plotted.

### Transmission Electron Microscopy

Transmission electron microscopy was performed as previously described (Galletta et al., 2020). Adult testes were fixed in 2.5% glutaraldehyde and 1% formaldehyde in 0.12 M sodium cacodylate buffer (pH 7.2) for one hour. Samples were post fixed in 1% osmium tetroxide in cacodylate buffer, stained with 1% uranyl acetate, then dehydrated in an ethanol series with propylene oxide. Samples were then embedded in Embed 812 resin (Electron Microscopy Sciences, Hatfield, PA, USA). 50-60 nm sections were obtained in transverse and longitudinal orientations using an EM UC7 ultramicrotome (Leica, Vienna, AUT). Sections were post-stained with uranyl acetate and lead citrate. Images were acquired on a JEM-1200EX (JEOL, USA) TEM with an accelerating voltage of 80 keV and equipped with an AMT 6 megapixel digital camera (Advanced Microscopy Techniques Corp) or a JEM 1400 (JEOL USA) TEM with an accelerating voltage of 120 keV and equipped with an AMT XR-111 digital camera (Advanced Microscopy Techniques Corp).

### DCTN3, Lis-1, and Nup358 quantification

For DCTN3, Lis-1, and Nup358 quantification, only early Round and Round stage spermatids were selected for analysis. Spag4 (from a *spag4::6myc* transgene) was used to ensure that all assessed nuclei were spermatid nuclei for DCTN3 and Lis-1 quantification. A line scan of the DCTN3, Lis-1, or Nup358 signal was taken by drawing a 10-pixel-wide by 2-um-long line perpendicular to the edge of the nucleus and centered on the nuclear edge in FIJI software. For spermatids with an intact HTCA, the line was drawn directly adjacent to the centriole. For spermatids without an intact HTCA, the line was drawn towards the mitochondrial derivative or towards the unattached centriole. Background signal adjacent to the nuclear envelope was eliminated by subtracting the average of the first five and last five values of the line scan for each spermatid. Intensity values were then plotted as a function of distance and the area under the curve was calculated in GraphPad Prism. Only peaks that reached at least 25% of the maximum value were used in the area calculation. Data was then normalized to the mean of the control for each experiment. For Lis-1 specifically, only testes in which some amount of Lis-1 signal was detectable were used for analysis, which likely leads to an underestimate of the severity of the phenotype.

### Statistics

For graphs with comparisons between two groups, an unpaired two-tailed t-test was performed. For graphs with comparisons between three or more groups, a one-way ANOVA with Tukey’s correction was used. Sample sizes are denoted in the figure legends. All statistical analysis was performed in GraphPad Prism 10.4.1 (GraphPad Software, Boston, MA, USA). Error bars represent the mean ± standard deviation for each graph. Colors represent different experimental trials.

---

**Supplemental Figure 1:**
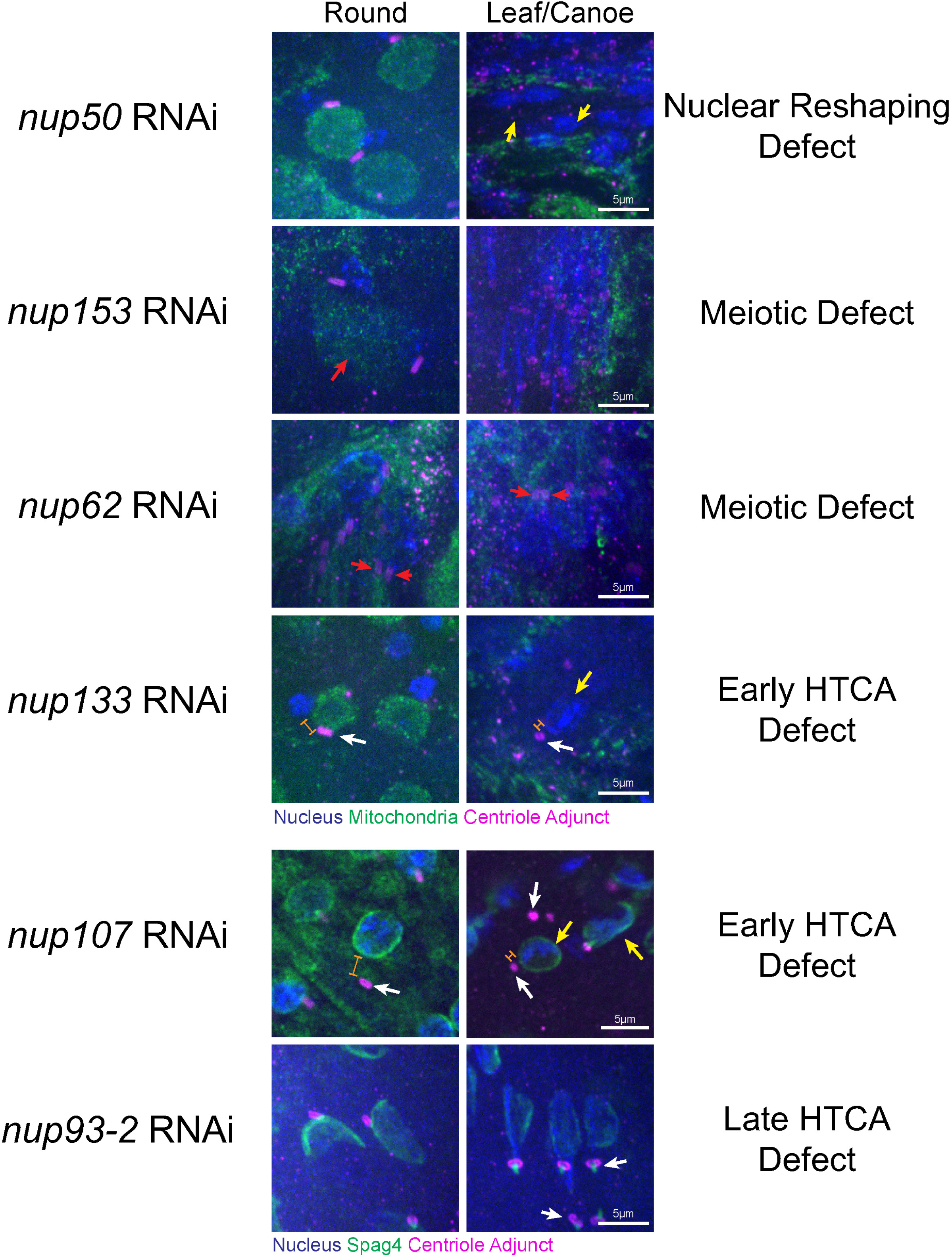
Nucleoporin RNAi screen revealed different phenotypes in spermiogenesis. Representative images showing the different phenotypes observed in the nucleoporin RNAi screen. All RNAi was driven by the testis-specific *Bam-Gal4* driver. Blue, nucleus (DAPI); magenta, centriole adjunct (Asl); green, mitochondria (ATP5A) or Spag4 (Spag4::6myc). Yellow arrows indicate nuclear shaping defects. Red arrows indicate meiotic defects with more than one centriole or nucleus in a single spermatid. White arrows indicate HTCA defects. Orange brackets indicate separation between nucleus and centriole. Scale bars 5um.

**Supplemental Figure 2:**
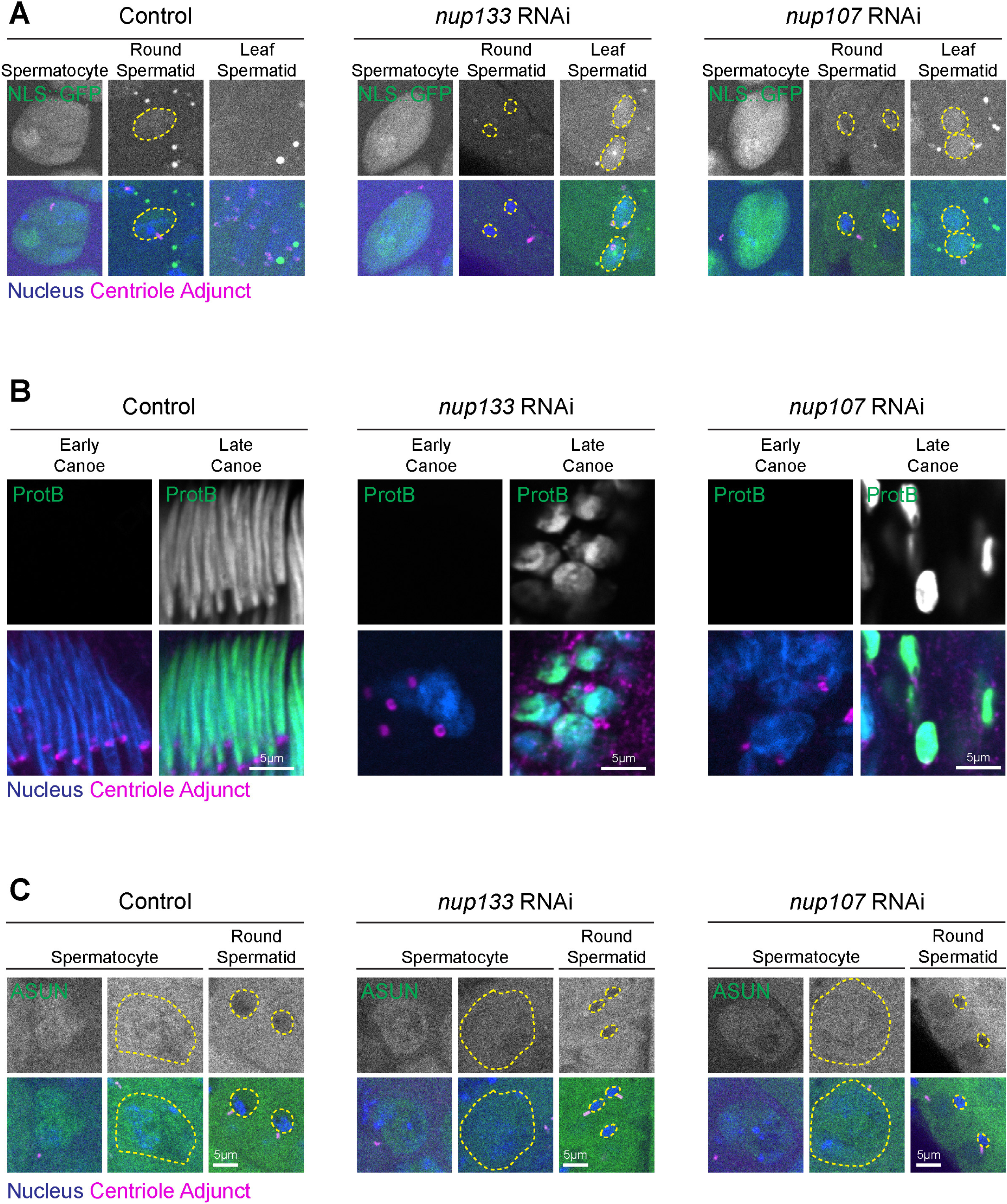
Nucleocytoplasmic transport still occurs following Nup depletion. **(A)** Representative images of Spermatocytes and Round and Canoe spermatids for the indicated genotypes. Blue, nucleus (DAPI); magenta, centriole adjunct (Asl); green, NLS::GFP. Dashed yellow line shows outline of nucleus. Scale bar 5um. **(B)** Representative images of Early Canoe and Late Canoe spermatids for the indicated genotypes. Blue, nucleus (DAPI); magenta, centriole adjunct (Asl); green, Protamine B (ProtB::GFP). Scale bar 5um. **(C)** Representative images of Spermatocytes and Round spermatids for the indicated genotypes. Blue, nucleus (DAPI); magenta, centriole adjunct (Asl); green, asunder (ASUN::GFP). Dashed yellow line shows outline of nucleus. Scale bar 5um.

**Supplemental Figure 3:**
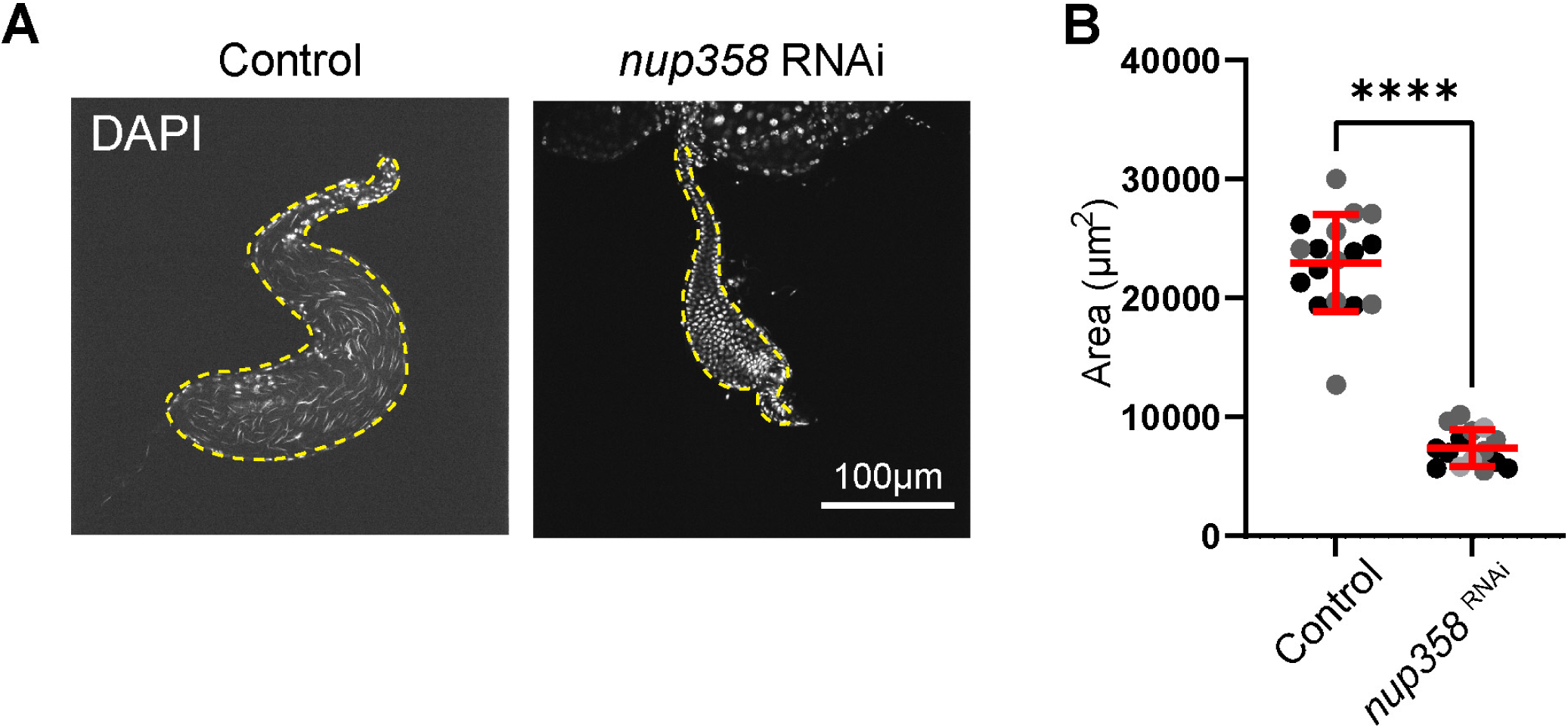
Nup358 is required for proper sperm production. **(A)** Representative images of seminal vesicles of the indicated RNAi knockdowns stained with DAPI (DNA). Yellow dashed outline shows seminal vesicles. Scale bar 100um. **(B)** Quantification of seminal vesicle area for controls (*Bam-Gal4/+,* n=17 seminal vesicles) and *nup358* RNAi (*nup358 RNAi/+; Bam-Gal4/+,* n=15 seminal vesicles); ****=*p*≤0.0001.

**Table S1:** Nucleoporin RNAi screen of HTCA phenotypes.

| **Nup** | **Type** | **RNAi line** | **Bam-Gal4 driven phenotype** |
| --- | --- | --- | --- |
| Nup50 | Nuclear basket | #34580<br>(Bloomington) | Impaired nuclear shaping,<br>possible HTCA defect |
| Nup153 | Nuclear basket | #30504<br>(Bloomington) | Meiotic defect |
| Mgtor (tpr) | Nuclear basket | #32941<br>(Bloomington) | Normal |
| Ndc1 | Transmembrane ring | #67275<br>(Bloomington) | Multiple centrioles; impaired<br>nuclear shaping |
| Gp210 | Transmembrane ring | #65219<br>(Bloomington) | Normal |
| Nup54 | Central channel | #57426<br>(Bloomington) | Normal |
| Nup58 | Central channel | #60110<br>(Bloomington) | Impaired nuclear shaping |
| Nup62 | Central channel | #35695<br>(Bloomington) | Meiotic defect; multiple<br>centrioles |
| Nup98-96 | Central channel | #28562<br>(Bloomington) | Normal |
| Nup93-2 | Inner ring | #51758<br>(Bloomington) | Late HTCA defect; impaired<br>nuclear shaping |
| Nup35 | Inner ring | #66002<br>(Bloomington) | Normal |
| Nup154<br>(Nup155) | Inner ring | #34710<br>(Bloomington) | Severe meiotic defect; HTCA<br>defect |
| Nup205 | Inner ring | #28610 (VDRC) | Meiotic defect/multiple<br>centrioles; HTCA defect;<br>impaired nuclear shaping |
| Nup188 | Inner ring | #102650 (VDRC) | Normal |
| **Nup133** | **Y-complex** | **#58290**<br>**(Bloomington)** | **Early HTCA defect; impaired**<br>**nuclear shaping** |
|  |  | **#110194 (VDRC)** | **Early HTCA defect; impaired**<br>**nuclear shaping** |
| Nup37 | Y-complex | #62328<br>(Bloomington) | Normal |
| Nup75 | Y-complex | #28315<br>(Bloomington) | Normal |
| Nup160 | Y-complex | #32391<br>(Bloomington) | Normal |
| Nup44A (Seh1) | Y-complex | #32942<br>(Bloomington) | Normal |
| Sec13 | Y-complex | #32468<br>(Bloomington) | Normal |
| Nup43 | Y-complex | #33645 (VDRC) | Meiotic defect; late HTCA<br>defect; impaired nuclear<br>shaping |
|  |  | #108595 (VDRC) | Severe meiotic defect; HTCA<br>defect |
| **Nup107** | **Y-complex** | **#22407 (VDRC)** | **Early HTCA defect; impaired nuclear shaping** |
|  |  | **#110759 (VDRC)** | **Early HTCA defect; impaired nuclear shaping; minor meiotic defect** |
| ELYS | Y-complex | N/A | N/A |
| Mbo (Nup88) | Cytoplasmic filaments | #77374 (Bloomington) | Multiple centrioles/meiotic defect |
| Nup214 | Cytoplasmic filaments | #33897 (Bloomington) | Meiotic defect/multiple centrioles; HTCA defect |
| Nup358 | Cytoplasmic filaments | #34967 (Bloomington) | No spermatocytes |
|  |  | #38581 (VDRC) | Severe meiotic defect; HTCA defect |
|  |  | #38583 (VDRC) | Normal |
| CG18787 (NLP1) | Cytoplasmic filaments | #55375 (Bloomington) | Normal |
| Rae1 | Peripheral | #32882 (Bloomington) | Severe meiotic defect; impaired nuclear shaping; HTCA defect |

